# Distribution models predict climate-related range alteration or extinction of eleven threatened tropical rainforest trees in the Western Ghats

**DOI:** 10.1101/2024.06.19.599808

**Authors:** A. P. Madhavan, Kshama Bhat, Srinivasan Kasinathan, Divya Mudappa, Navendu Page, T. R. Shankar Raman

## Abstract

Evidence for climate-change related alteration in distributions and ranges of forest trees is accumulating, but information from Asian tropical forests, particularly for threatened and endemic species, remains limited. Here, we examine landscape-level distribution-abundance patterns of 11 endemic and threatened tropical rainforest tree species in the Anamalai Hills and model their distribution and responses to future climate scenarios in the Southern Western Ghats (SWG), India. Six of the primarily low- and mid-elevation species were more abundant in protected reserves than forest fragments. Occurrence data from the Anamalais and SWG (*N =* 3004, range: 41 – 706 per species) were used to model distributions using maximum entropy (maxent) species distribution modelling in R. Maxent model performances indicated excellent fits for 9 of 11 species (AUC>0.90) with precipitation and temperature variables showing higher permutation importance. There was high interspecies variability in range size (197 – 12,221 km^2^) and niche width (0.04 – 0.50). Models of distribution under future climate in 2061 – 2080 predict range reductions in six species (including near-extinction for two species), increases for three species, and no substantial change for two species. Predicted southward and westward shifts in ranges and persistence in parts of the SWG indicate the importance of identifying and conserving micro-climatic zones or refugia to ensure the persistence of tree species under anticipated climate change.

## INTRODUCTION

Global climate change continues to have ongoing and accelerating effects on broad-scale ecosystem processes, population dynamics, and phenology of species, besides directly influencing their distribution patterns (Scheffers et al., 2016). Climate-influenced distribution patterns are linked to the ecophysiological tolerances, thresholds and sensitivities of species to prevailing climate (Davis & Shaw, 2001; Woodward & Williams, 1987). Disequilibrium and shifts in prevailing environmental conditions due to climate change may positively or negatively influence the growth rates and persistence of species with the magnitude of change depending on their climatic sensitivities (Charney et al., 2016). To persist under future climate change, species will be forced to adapt, migrate, or face extirpation (Aitken et al., 2008; Davis & Shaw, 2001). Such migrations and shifts (contraction/expansion) in species distribution patterns (Roberts & Hamann, 2016) may occur under varying velocities (Loarie et al., 2009) towards associated suitable or stable climate refugia characterised by low climatic fluctuations (Gavin et al., 2014; Stebbins & Major, 1965). Climate refugia act as cradles for endemism and species diversification (Gopal et al., 2023; Harrison & Noss, 2017) and understanding how species occurrence and distribution patterns change with environmental and climate gradients can throw light on future adaptation or extirpation scenarios in these regions (Bonachela et al., 2021; Neilson et al., 2005).

Within India, the southern latitudes of the Western Ghats mountain range have been characterised as ecologically-significant climate refugia (Bose et al., 2019; Joshi & Karanth, 2013). The southern Western Ghats (SWG) sustained ecological diversity through significant post-glacial warming and cretaceous volcanism of the Deccan Plateau, enabling species distribution expansions and successive migrations (Joshi & Karanth, 2013). The topography of this mountain range, its north-south axis, and parallel orientation to the coast, have resulted in distinct present-day latitudinal and elevational gradients (Davidar et al., 2007). Lower latitudes present more stable climates with lower seasonality, water deficit, and temperature fluctuations, while also having greater topographic complexity and taller mountains. Along the elevational gradient, higher elevations have lower temperatures, while a moisture gradient exists from wetter western slopes to drier eastern rainshadow regions (Krishnadas et al., 2016; Page & Shanker, 2020; Pascal, 1988). Environmental gradients manifest across different spatial scales with macro-scale (latitudinal) variation in temperature and precipitation seasonality and local mesoscale (elevational) variation in precipitation and temperature thresholds (Cayuela et al., 2006; Krishnadas et al., 2016; Pascal, 1988). Under these gradients, compared to the northern regions, the southern Western Ghats tree species assemblages are more diverse and specialised (Davidar et al., 2007; Gopal et al., 2023; Krishnadas et al., 2021; Page & Shanker, 2020) with higher rates of endemism and species turnover (Bose et al., 2019; Ramesh & Pascal, 1997). More than half the tree species within this zone of climatic stability are endemic to the southern region (Bossuyt et al., 2004; Gunawardene et al., 2007).

Past research on trees and woody plants in the SWG has attested the region’s phylogenetic importance (Bose et al., 2019; Chitale et al., 2014; Gopal et al., 2023; Joshi & Karanth, 2013) and documented patterns in plant community composition and diversity across environmental gradients (Davidar et al., 2007; Krishnadas et al., 2016, 2021; Page & Shanker, 2020; Pascal, 1988; Vijayakumar et al., 2016). The role of the SWG as climate refugia under future climate change and anticipated changes in species distributions are as yet poorly documented. Previous species distributional studies have primarily focused on single tree species or specific families (Deb et al., 2017; Malik et al., 2022; Pramanik et al., 2018; Priti et al., 2016; Shivaprakash et al., 2022) and deriving generalities has remained a challenge due to varying methods, sample sizes, and types of models used. Further, there has been limited attention on threatened and endemic tree species that may be particularly vulnerable to the effects of climate change besides historical and ongoing landscape alteration and habitat fragmentation (Muthuramkumar et al., 2006; Reddy et al., 2016).

By using, as in the present study, robust species distribution models on a selection of threatened tree species with varying climate sensitivities, thresholds, and distributions, one can better determine patterns in individual species responses to climate change (Godoy-Veiga et al., 2021) and potential future trajectories of species groups that share specific affinities (Aitken et al., 2008). Here, we explore species distribution models of 11 threatened tree species in 11 genera, combining information from published sources with new distributional surveys conducted in the Anamalai Hills in the southern Western Ghats. Our main objectives were: (a) to first examine landscape-level distribution and abundance patterns of the focal species in relation to elevation and habitat fragmentation; (b) to examine relationships between climatic variables and species distribution to develop species-specific spatial distribution models and estimate distributional ranges; and (c) to use the developed models to assess species-specific impacts of future climate change projections on distributional ranges in the southern Western Ghats. The results help identify patterns in species responses in relation to climate sensitivities and highlight the importance of climatic refugia for tree species in the context of projected climate change in tropical forest regions.

## MATERIALS AND METHODS

### Study species

The study focused on 11 tree species in 11 genera and 10 families placed under various threat categories in the *IUCN Red List* (IUCN, 2022). This includes 2 Critically Endangered (*Dipterocarpus bourdillonii, Phyllanthus anamalayanus*), 4 Endangered (*Cryptocarya anamalayana*, *Dysoxylum malabaricum*, *Orophea thomsonii*, *Palaquium ravii*), and 5 Vulnerable (*Diospyros paniculata*, *Drypetes wightii*, *Myristica beddomei*, *Syzygium densiflorum*, *Vateria indica*) species. All 11 species are primarily found in tropical moist forests (tropical moist deciduous to wet evergreen forests) in the Western Ghats (Ramesh & Pascal, 1997). Barring four species (*D. paniculata*, *M. beddomei*, *V. indica, and Dysoxylum malabaricum*) that were more widely distributed, the remaining species were restricted to the southern Western Ghats (SWG, south of 13°N).

### Study area

The Western Ghats mountain range along India’s west coast is a global biodiversity hotspot (Kumar et al., 2004). Tropical moist forests in the Southern Western Ghats (SWG), which were the focus of this study, are encompassed within two terrestrial ecoregions (Wikramanayake et al., 2002): Southern Western Ghats Montane Rain Forests, Southern Western Ghats Moist Deciduous Forests. The SWG (73.95° – 80.33° E, 8.06° – 13.11°N) extends for 600 km along the southwestern coastline of the Indian peninsula and ranges in elevation from 40 m to 2695 m. This broad elevation range encompasses distinct floristic communities (Champion & Seth, 1968; Pascal et al., 2004) including: low-elevation tropical wet evergreen and moist deciduous forests (mostly below 800 m), mid-elevation tropical wet evergreen forests (800 – 1450 m asl), and higher elevation tropical wet evergreen forests and shola-grassland mosaics (>1450 m asl). The mid-elevation tropical wet evergreen forests were the primary vegetation type for the majority of the study species. The mean annual precipitation is around 2800 mm occurring primarily during the southwest monsoon (June – August).

The intensive study site in the Anamalai Hills of the SWG encompassed the moist western tracts of the Anamalai Tiger Reserve (ATR), Tamil Nadu, India (core zone: 958 km², 10.2160N, 76.8160E – 10.5660N, 77.4160E) and the adjoining Valparai Plateau (220 km², 10.250N, 76.8660E – 10.3660N, 76.9830E). Fieldwork in ATR was concentrated in the Valparai, Manamboli, and Ulandy Ranges that contain most of the remaining tropical wet evergreen forests, spanning elevations between 600 m and 1950 m. Tea and coffee plantations dominate the Valparai Plateau which also contains over 50 rainforest fragments (1 ha to >300 ha) embedded within the production landscape (Muthuramkumar et al. 2006; Mudappa & Raman 2007). The natural vegetation type falls mainly within the mid-elevation (700–1,400 m) tropical wet evergreen forest of the *Cullenia exarillata* – *Mesua ferrea* – *Palaquium ellipticum* type (Pascal 1988).

### Occurrence data

Occurrence data were gathered using a combination of intensive field surveys in the Anamalai Hills of the southern Western Ghats (Figure 1) and compilation of distributional records over the wider study region from earlier literature and published sources.

**Figure 1:**
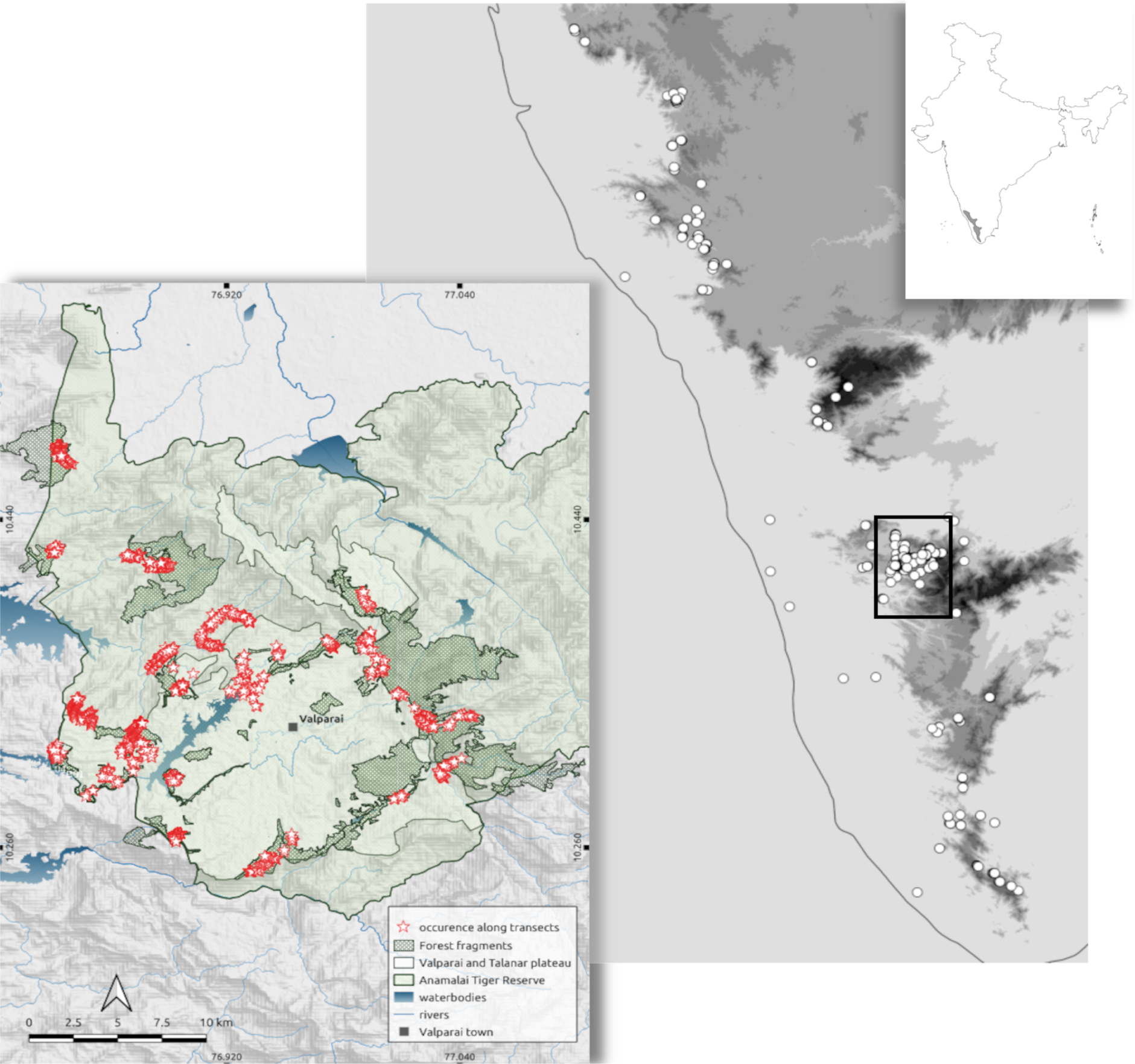
Map showing location of the Southern Western Ghats (SWG) in India (inset on top right), the SWG region with illustrative occurrences (centre), and occurrences of focal species within the intensive study site of the Anamalai Hills (on left).

During (2021-22), geographic occurrence locations of 11 threatened trees were collected from the Anamalai hill range (nested within the SWG complex). Intensive field surveys were carried out in 29 sites along 65 trails of 113.35 km total length in rainforest fragments of the Valparai Plateau (VP) and mature tropical rainforests within the Anamalai Tiger Reserve (ATR). This included 16 rainforest fragment sites in VP (26 trails, 35.46 km, 800 m to 1400 m) and 13 rainforest sites in ATR (39 trails, 77.89 km, 580 m to 1940 m). Trails were walked to cover as much of the area within fragments in VP as well as cover different locations and altitude zones (between 600 m and 2000 m) in ATR. Along each trail, we recorded occurrences of the focal 11 threatened tree species within 10 m on either side of these trails, noting species identity and characteristics such as location, elevation, girth at breast height, height, canopy shape, damage, and other attributes.

We combined the distribution data from the Anamalai hills (1940 individuals) with occurrence records downloaded from the Global Biodiversity Information Facility (GBIF, https://www.gbif.org) and extensive earlier surveys between 2010 – 2014 by (Page & Shanker, 2020), yielding a total of 3004 records (11 species) distributed across the Southern Western Ghats. This combined dataset provides a reasonable representation of the variation in environmental conditions, latitudinal and altitudinal extents, and geographic clustering of the focal species.

### Analysis

In the ATR landscape, we used all occurrences to examine spatial and elevational distribution of species. Species densities were calculated along all trails longer then 0.2 km (*N* = 60) from the total number of individuals recorded divided by surveyed area (trail length x 20 m).

### Environmental variables

Analysis was conducted using the Bioclim environmental data (resolution 1 km^2^) from the Worldclim raster database (Fick & Hijmans, 2017). These raster files were cropped to the latitudinal extent of the southern Western Ghats montane rainforest ecoregion bounds/extents. We used variation inflation factor analysis to assess high collinearity of the 19 Bioclim variables and select a subset of 7 variables with reduced collinearity. The 7 variables subsequently used in all fitted models were: precipitation of the driest month (pdm), mean temperature of wettest quarter (mtwq), mean diurnal range (mdr), precipitation of warmest quarter (pwaQ), precipitation of coldest quarter (pcQ), precipitation seasonality (PS, coefficient of variation).

### Species distribution modelling

Analyses were carried out within the R statistical and programming environment (R Core Team, 2023) using the ‘flexsdm’ package (Velazco et al., 2022). The package ‘flexsdm’ offers refined functionality to account for spatial and environmental biases due to sampling intensity, and geographic and environmental clustering (Phillips et al., 2009). The analyses used the ‘terra’ package (Hijmans et al., 2023) for manipulation of spatial raster data and the ‘SDMtune’ package for post-modelling applications, e.g., assessing variable importance (Vignali et al., 2020).

We sampled pseudo-absence and background points within a buffer of radius 20 km, 30 km, or 40 km around the occurrence points, with smaller buffers for more range-restricted species. To sample pseudo-absence and background points, a spatial block partitioning method was used (Roberts et al., 2017). The method spatially divided the occurrence locations into grids (some idea about the size of the grids would be useful). Each of these grids is then assigned to one of the 2, 3 or 4 partitions, depending on the spatial distribution and number of occurrence locations. These grids served as spatial units for sampling background and pseudo-absence points besides also functioning as a method for partitioning occurrence locations into training and testing data). The optimum grid size for each species was selected through inbuilt environmental and spatial autocorrelation methods testing multiple inputs and finding the best fit maximising representation of occurrence points between the grids and their partitions. Background points were randomly generated within each spatial partition (Vollering et al., 2019) with the total number of background points being ten times the number of occurrences in each partition. Pseudo-absence points, equal in number to occurrences in each spatial partition, were generated using environmental constraints avoiding overlap with the environmental conditions i.e. cells with exactly the same values of seven environmental variables as that of the occurrence locations were avoided (Barbet-Massin et al., 2012). For species with larger sample sizes, we used both environmental and geographic constraints (an added minimum distance from presences to sample absence points) to avoid overlap with occurrence locations.

### Modelling procedures

Maxent species distribution models (Phillips et al., 2006) were chosen for species distribution modelling of all 11 species using the seven selected environmental variables. Maxent models were chosen after comparing performance parameters across Maxent, generalised linear models, neural network machine learning models, gaussian models, boosted regression models, and ensemble model approaches. Performance parameters considered included area under the Receiver Operating Characteristics curve (AUC, with AUC≥0.7 taken to indicate reasonable fit, ≥0.8 indicating good fit, and ≥0.9 indicating excellent fit), Total Sum of Squares (TSS), True Positive Rate (TPR), True Negative Rate (TNR), and Sorensen index. Maxent models had the highest performance scores for this dataset, and also appeared to generate the most reasonable species distribution predictions. Models were tuned using linear-quadratic and linear-quadratic-hinge based classes chosen based on inbuilt model evaluations of the highest scores. Two threshold types were considered including ‘max_sens_spec,’ the threshold at which the sum of the sensitivity and specificity is the highest, and ‘equal_sens_spec,’ the threshold at which sensitivity and specificity are equal (Merow et al., 2013). Regularisation values were selected by the Maxent models from a standardised sequence of values. Occurrence-based constraints were imposed to prevent model overfitting (Mendes et al., 2020). The threshold value for the posterior constraints were based on the tuned maxent models.

The fitted Maxent model was then used to predict occurrence probabilities across the southern Western Ghats study area. The permutation importance (100 iterations) of the seven variables was calculated for each species based on relative changes in AUC scores (normalised to percentages). The permutation importance values (strength of coefficients) together with climate response curves were taken to illustrate species niche generality/specificity and the nature of relationships to the environmental variables. Niche breadth was quantified using Levins’ B2^geo^ metric for niche speciality (Evans & Jacquemyn, 2022) using the functionality of the ‘ENMTools’ R package (Warren et al., 2021).

### Future projections

Using the same seven Bioclim variables, Maxent species niche models were projected to future global climatic scenarios chosen from among the Coupled Model Intercomparison Project Phase (CMIP6) models under the Sixth Assessment Report (AR6) of the Intergovernmental Panel on Climate Change (IPCC, 2022). We include all climate scenarios from optimistic to worst-case Shared Socio-economic Pathways (SSPs) risk assessments: SSP1-2.6, SSP2-4.5, SSP3-7.0, and SSP5-8.5. We considered SSP2-4.5 as the ‘high-likelihood’ climate change scenario (Hausfather & Peters, 2020). Out of the 36 global climatic models (GCMs) listed under CMIP6, the MIROC6 model (Kataoka et al., 2020; Tatebe et al., 2019) was selected for the time period of 2061-2080 based on the relative accuracy of this model to map the Indian monsoon over the last 50 years (Katzenberger et al., 2021). We estimated species geographic range extents’ as the estimated occurrence/projected area (km^2^) occupied by cells with a Gibbs value greater ≥0.3. Current and future species range extents were then compared using percentage change in area and visualised directionality and distributional change on maps.

## RESULTS

### Distribution and density at the landscape level

On field surveys in the Anamalai Hills, we recorded 1543 individuals of all the 11 focal tree species in the Anamalai Tiger Reserve (ATR, ‘Protected Area’) and 397 individuals of 10 focal species in rainforest fragments on the Valparai Plateau (VP, ‘Fragments’), making a total of 1940 individuals across the 65 routes. Of the 65 routes, *Dipterocarpus bourdillonii* occurred in 5 routes (7.7%) while *Myristica beddomei* occurred in 42 routes (64.6%, Table 1). Elevational distribution patterns indicated that the median elevation of occurrence was below 1000 m for six species (*D. bourdillonii, Vateria indica, Orophea thomsonii, Diospyros paniculata, Palaquium ravii*, *M. beddomei*) and was above 1000 m for five species (*Dysoxylum malabaricum, Cryptocarya anamalayana, Drypetes wightii, Phyllanthus anamalayanus, Syzygium densiflorum*). Within the elevational limits of the study area, the low-elevation rainforest species *D. bourdillonii* had the lowest median and narrowest elevational range while the mid- and high-elevation *S. densiflorum* had the highest median and widest elevational range (Figure 2).

**Figure 2:**
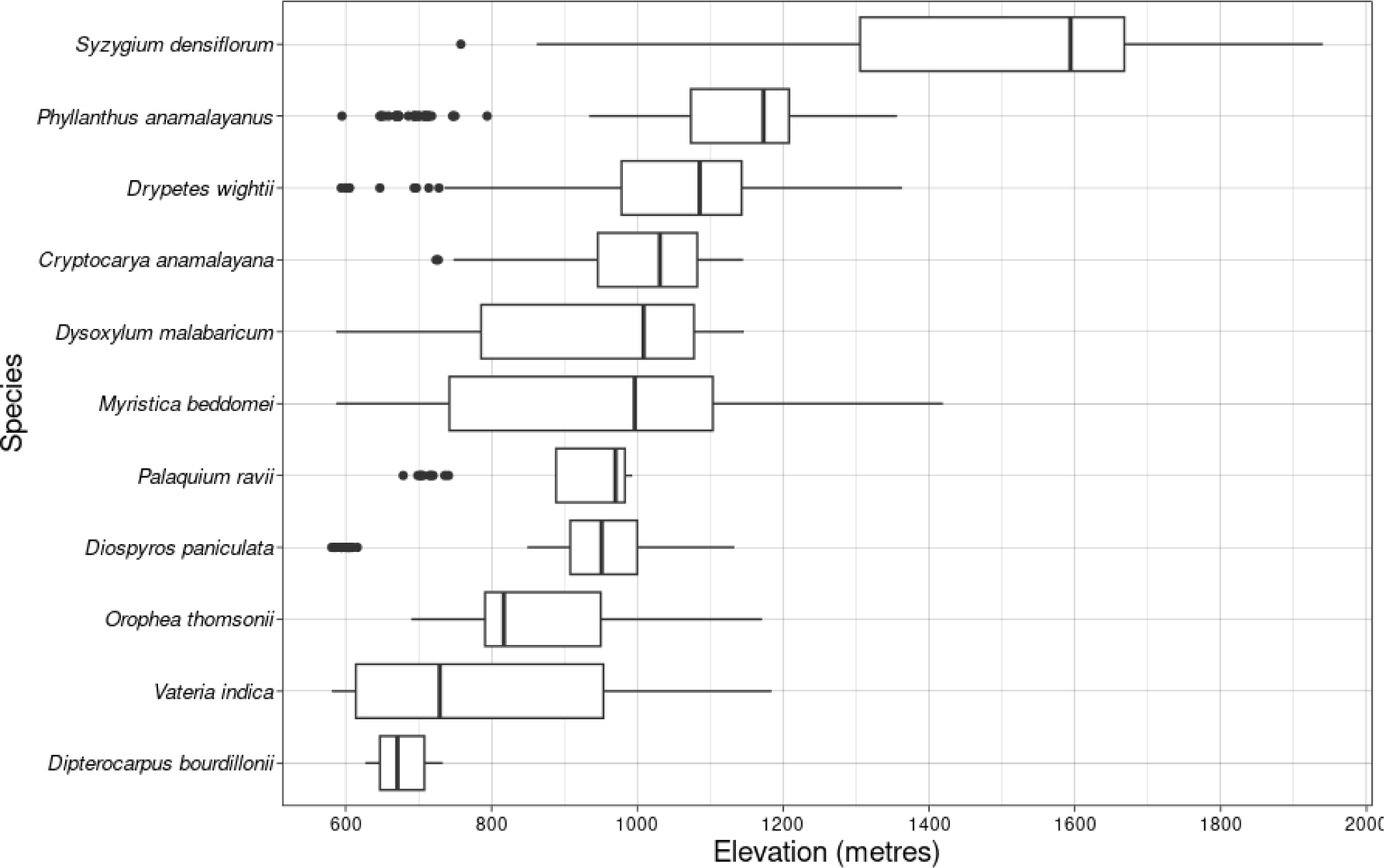
Boxplot of elevational distribution of the 11 threatened tree species in the Anamalai Hills, Southern Western Ghats, India

**Table 1:** Occurrence and distribution (percentage of routes where a species occurred) of focal threatened tree species in the Anamalai Hills, Southern Western Ghats.

| Species | Individual occurrences (routes) |  |  | Distribution (%) |  |  |
| --- | --- | --- | --- | --- | --- | --- |
|  | All | Protected area | Fragments | All | Protected area | Fragments |
| *Cryptocarya anamalayana* | 66 (16) | 44 (8) | 22 (8) | 24.6 | 20.5 | 30.8 |
| *Diospyros paniculata* | 117 (19) | 67 (8) | 50 (11) | 29.2 | 20.5 | 42.3 |
| *Dipterocarpus bourdillonii* | 37 (5) | 37 (5) | - (-) | 7.7 | 12.8 | - |
| *Drypetes wightii* | 278 (38) | 218 (23) | 60 (15) | 58.5 | 59.0 | 57.7 |
| *Dysoxylum malabaricum* | 142 (29) | 100 (15) | 42 (14) | 44.6 | 38.5 | 53.8 |
| *Myristica beddomei* | 280 (42) | 239 (26) | 41 (16) | 64.6 | 66.7 | 61.5 |
| *Orophea thomsonii* | 250 (20) | 229 (14) | 21 (6) | 30.8 | 35.9 | 23.1 |
| *Palaquium ravii* | 44 (7) | 42 (6) | 2 (1) | 10.8 | 15.4 | 3.8 |
| *Phyllanthus anamalayanus* | 183 (19) | 121 (15) | 62 (4) | 29.2 | 38.5 | 15.4 |
| *Syzygium densiflorum* | 152 (29) | 133 (18) | 19 (11) | 44.6 | 46.2 | 42.3 |
| *Vateria indica* | 391 (30) | 313 (18) | 78 (12) | 46.2 | 46.2 | 46.2 |

Density of species varied between Protected Area and Fragments (Figure 3) in all but three species (*C. anamalayana, D. malabaricum, P. anamalayanus*). Only one species (*D. paniculata*) occurred at significantly higher density in the Fragments stratum (Poisson GLM, *P* < 0.01). Of the seven remaining species, *D. bourdillonii* was absent in Fragments, and the other six occurred at significantly higher densities within the Protected Area (*P* < 0.001).

**Figure 3:**
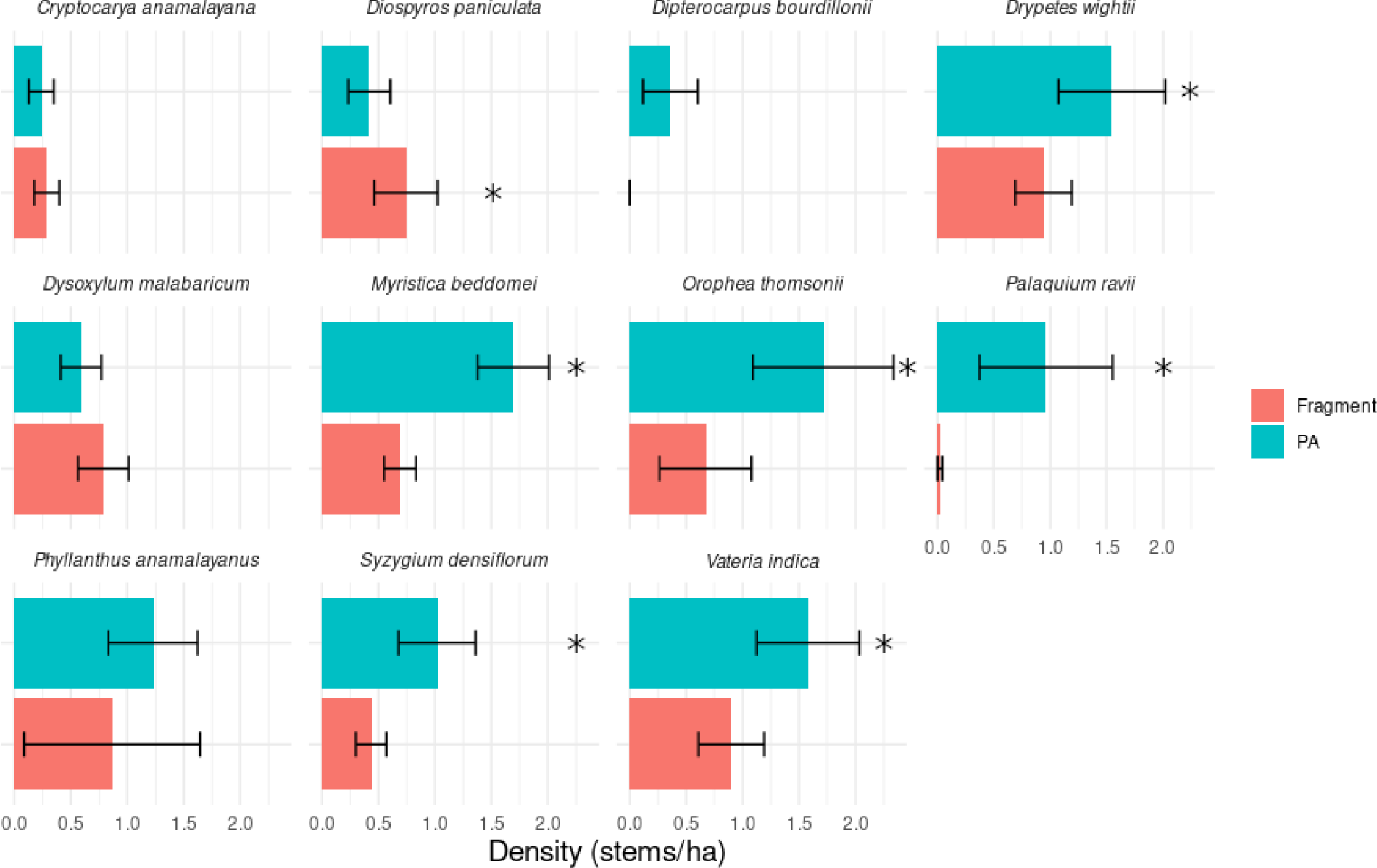
Differences in density of focal threatened tree species within the Anamalai Tiger Reserves (‘Protected Area’) and in rainforest fragments of the Valparai Plateau (‘Fragments’), Anamalai Hills, Southern Western Ghats. Statistical significance of differences (*P* < 0.01) as assessed using Poisson GLMs are indicated with an asterisk.

### Species distribution models: regional patterns

Species distributions across the Southern Western Ghats (SWG) region were modelled using Maxent models with the ‘flexsdm’ package. Maxent model performances indicated excellent fits for 9 of 11 species (AUC>0.90), good fit for *Diospyros paniculata* and *Vateria indica* (AUC=0.83; Table 2). True Positive Rate (TPR) was >0.9 for 10 species and 0.84 for *Cryptocarya anamalayana*, and True Negative Rate (TNR) was >0.9 for 9 species and >0.8 for *D. paniculata* and *V. indica*. Precipitation seasonality, precipitation of the warmest quarter, and precipitation of the driest month were identified as significant influences on species distributions of most species, followed by mean temperature of the wettest quarter and isothermality (Figure 4). There was considerable interspecific variation in which variables were more important. Variables with low overall influence (precipitation of the coldest quarter, mean diurnal range) were still important for a few species such as *Dysoxylum malabaricum*, *Syzygium densiflorum*, and *Dipterocarpus bourdillonii* (Figure 4).

**Figure 4:**
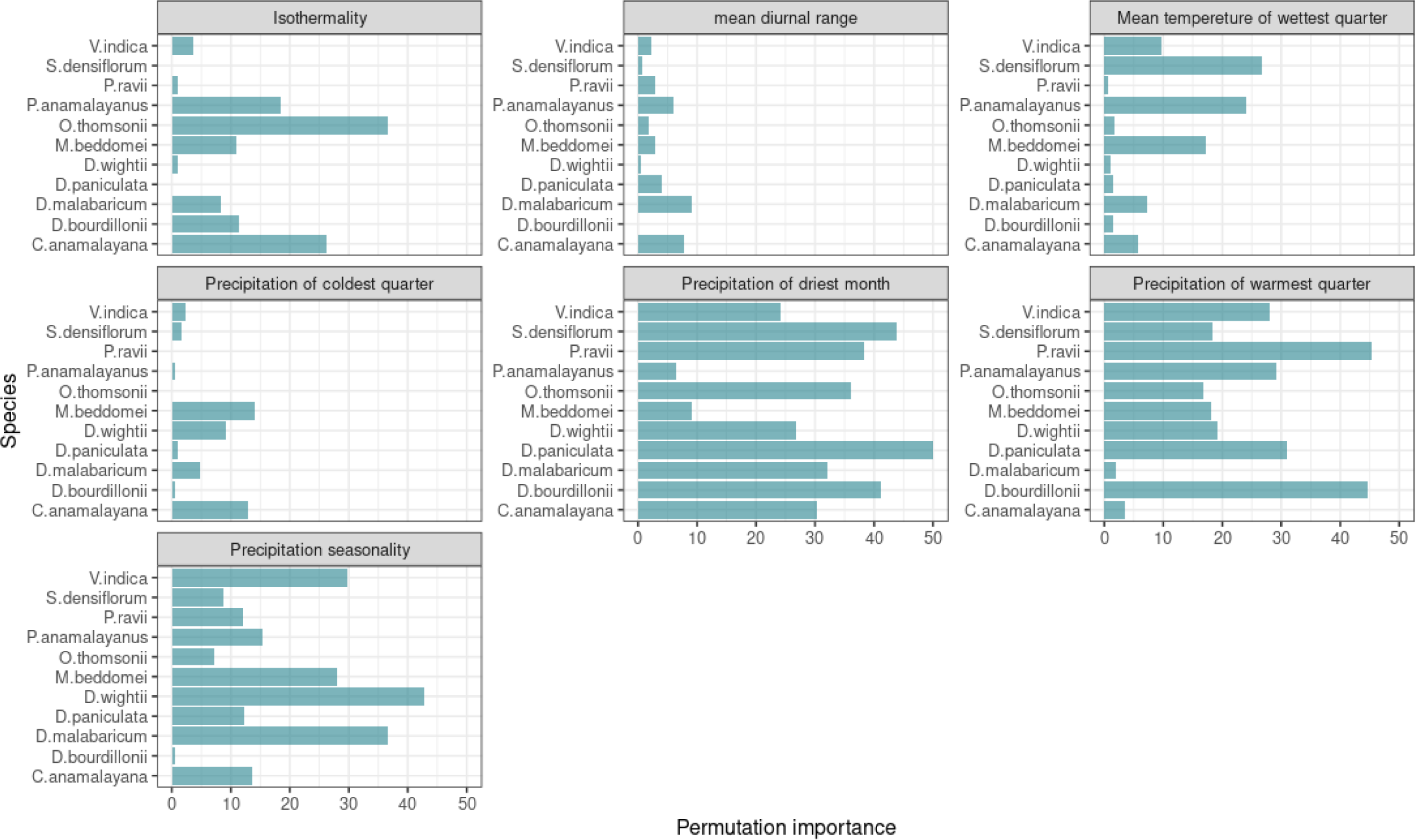
Permutation importance (strength of coefficients) of each environmental variable in influencing distributions for the 11 species.

**Table 2:**
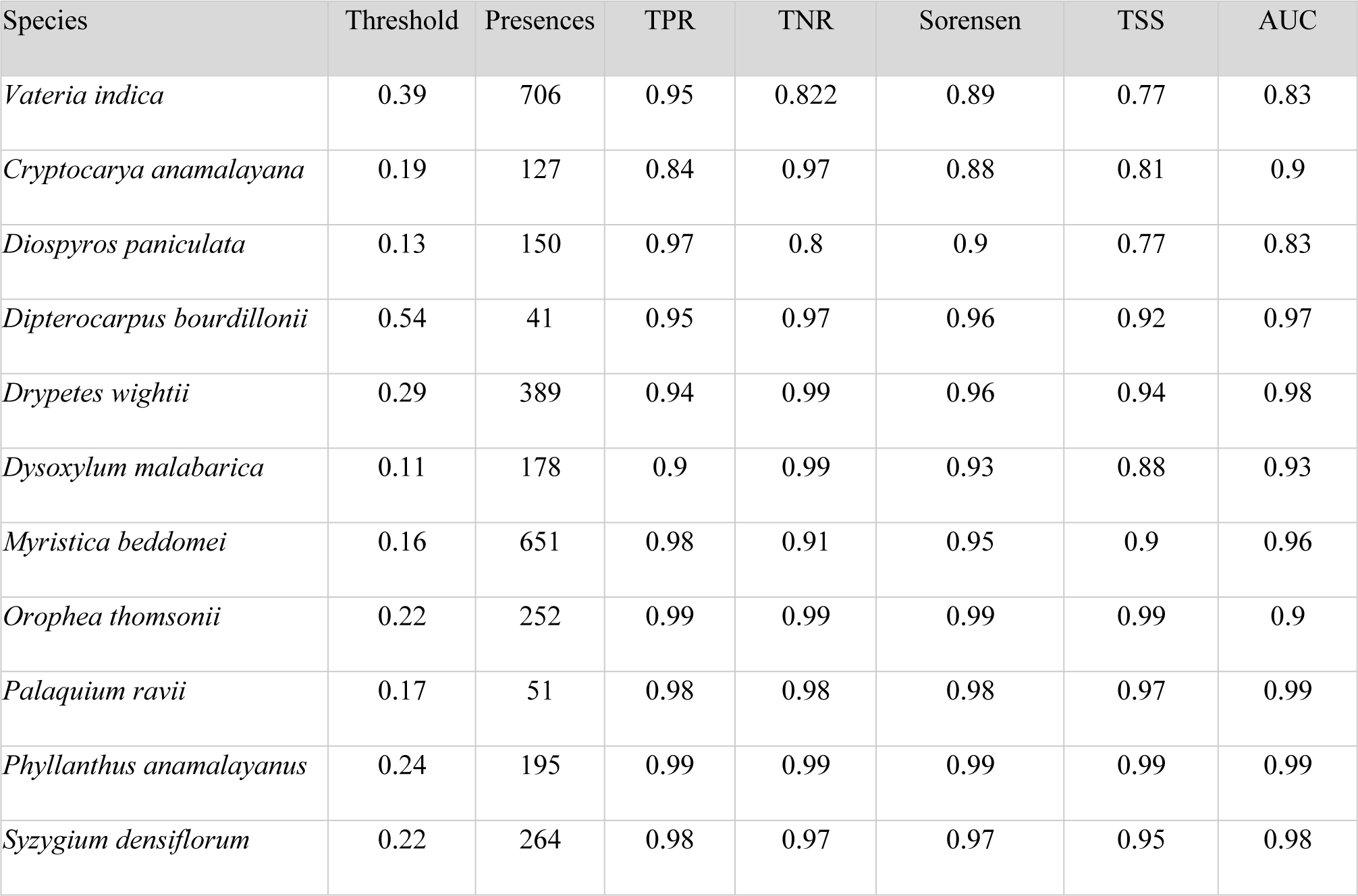
A comparison of model performance scores for the 11 species. Selected metrics include: true positive rate (TPR), true negative rate (TNR), Sorensen index mean, true skill statistic mean (TSS), and area under the receiver operating characteristics curve, mean (AUC).

| Species | Threshold | Presences | TPR | TNR | Sorensen | TSS | AUC |
| --- | --- | --- | --- | --- | --- | --- | --- |
| *Vateria indica* | 0.39 | 706 | 0.95 | 0.822 | 0.89 | 0.77 | 0.83 |
| *Cryptocarya anamalayana* | 0.19 | 127 | 0.84 | 0.97 | 0.88 | 0.81 | 0.9 |
| *Diospyros paniculata* | 0.13 | 150 | 0.97 | 0.8 | 0.9 | 0.77 | 0.83 |
| *Dipterocarpus bourdillonii* | 0.54 | 41 | 0.95 | 0.97 | 0.96 | 0.92 | 0.97 |
| *Drypetes wightii* | 0.29 | 389 | 0.94 | 0.99 | 0.96 | 0.94 | 0.98 |
| *Dysoxylum malabarica* | 0.11 | 178 | 0.9 | 0.99 | 0.93 | 0.88 | 0.93 |
| *Myristica beddomei* | 0.16 | 651 | 0.98 | 0.91 | 0.95 | 0.9 | 0.96 |
| *Orophea thomsonii* | 0.22 | 252 | 0.99 | 0.99 | 0.99 | 0.99 | 0.9 |
| *Palaquium ravii* | 0.17 | 51 | 0.98 | 0.98 | 0.98 | 0.97 | 0.99 |
| *Phyllanthus anamalayanus* | 0.24 | 195 | 0.99 | 0.99 | 0.99 | 0.99 | 0.99 |
| *Syzygium densiflorum* | 0.22 | 264 | 0.98 | 0.97 | 0.97 | 0.95 | 0.98 |

All 11 species had clear thresholds, tolerance limits, and sensitivities to climate gradients (Supplementary Information: Figure S1), corresponding to distinctive species distributional patterns under the Maxent models (Figure 5). Within the SWG, *Phyllanthus anamalayanus, Palaquium ravii, Orophea thomsonii,* and *Dipterocarpus bourdillonii* exhibited restricted distributions, compared to *Vateria indica, Diospyros paniculata,* and *Myristica beddomei* that showed broader distributions (Figure 5). *Cryptocarya anamalayana, Syzygium densiflorum, Drypetes wightii,* and *Dysoxylum malabaricum* were wider in distribution but only within the limits of the mid-latitudes of SWG (Figure 5).

**Figure 5:**
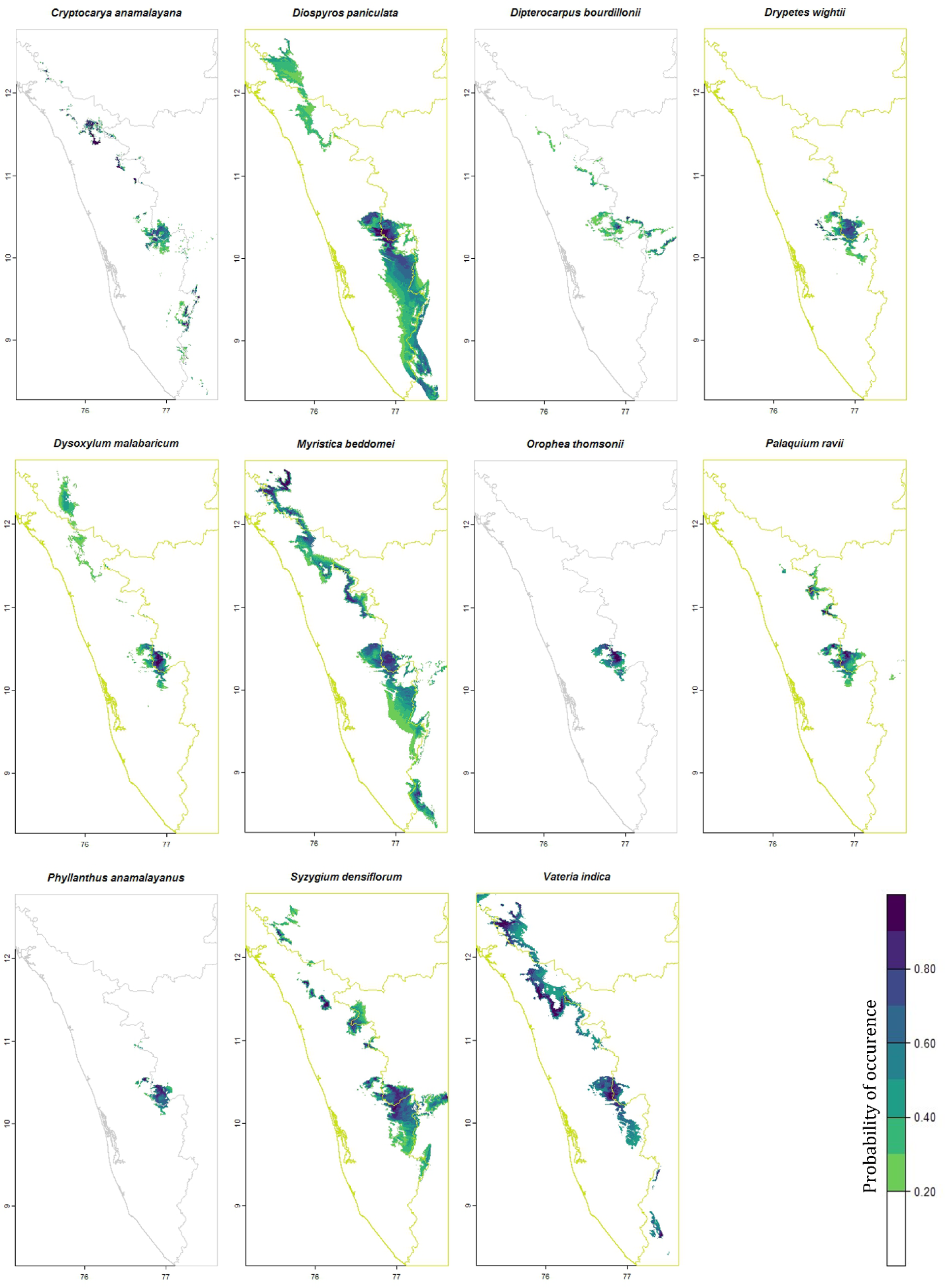
Species distributions of 11 threatened tree species in the southern Western Ghats derived from Maxent models. Darker colours indicate higher probability of occurrence.

### Distributional range and alteration under future climate scenarios

Considering areas with Gibbs value ≥0.3 as indicating species occurrence, the estimated distributional ranges show high variation across species, from 197 km² in *Dipterocarpus bourdillonii* to 12,221 km² in *Diospyros paniculata* (Figure 6). Niche breadth (Levins’ B2^geo^ values) showed a broad range from 0.04 in *Phyllanthus anamalayanus* to 0.5 in *Diospyros paniculata* (Supplementary Information: figure S2).

**Figure 6.**
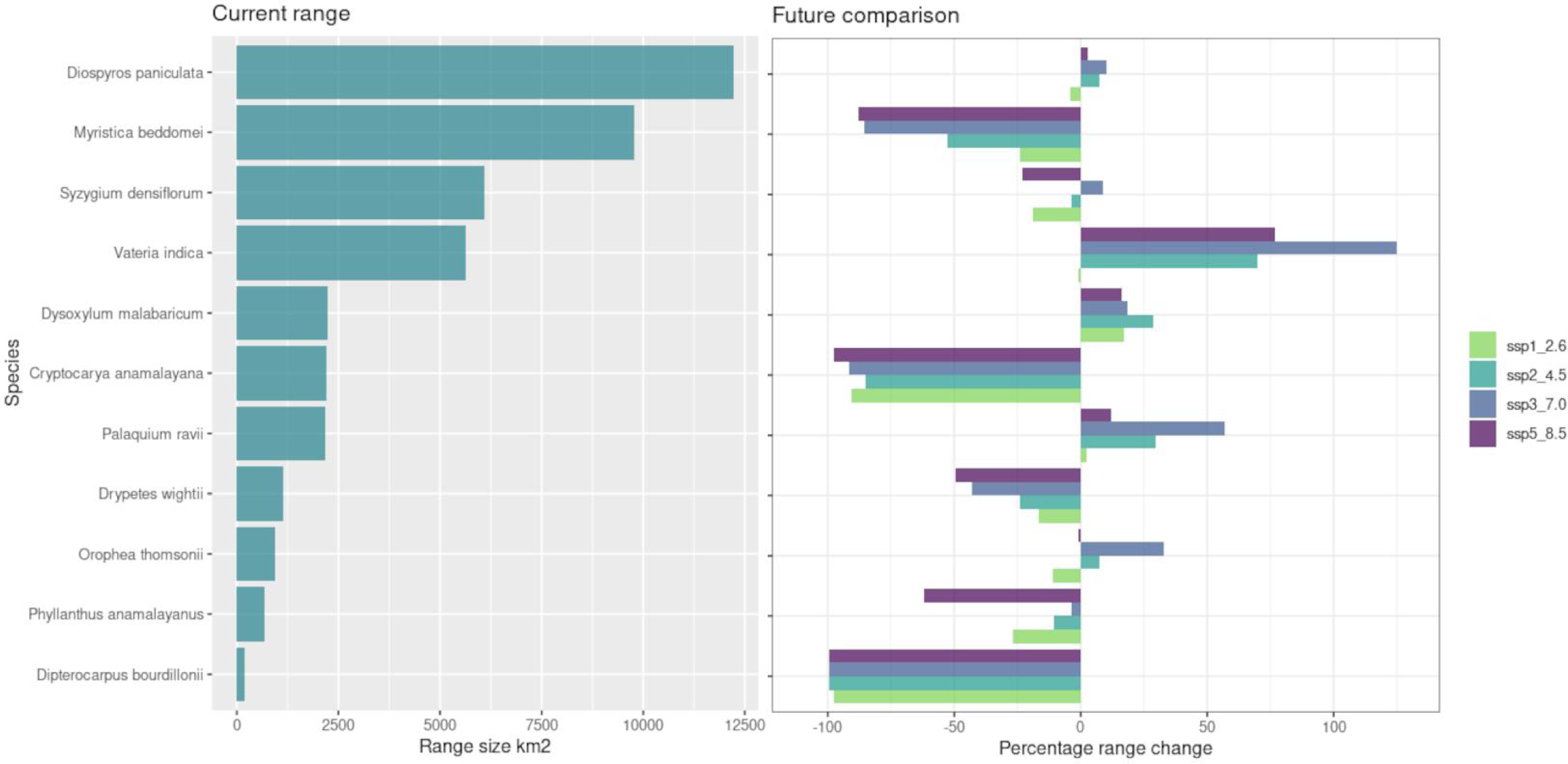
Left: Current estimates of climatically suitable range extent (area km^2^) for the 11 species within the SWG. Right: Relative percentage change in species’ distributional ranges in 2061 – 2080 under different future climate model scenarios.

Models for *Dipterocarpus bourdillonii* and *Cryptocarya anamalayana* predicted extinctions or near extinctions (>95% reduction in range) irrespective of climate change scenarios (Figure 6). For five species, models predicted progressively greater declines in range under more drastic climate change scenarios (SSP3-7.0 and SSP5-8.5) of future climate (2061-2080). For species that are predominantly distributed at low-elevation (below 1000 m elevation), models predicted range expansions across all scenarios for *Vateria indica* while *Dysoxylum malabaricum, Palaquium ravii* showed increases under intermediate SSP scenarios followed by drastic declines under SSP5-8.5. *Diospyros paniculata* showed little or no change under various scenarios. *Syzygium densiflorum, Orophea thomsonii* exhibited oscillating signals with minor reductions and increases in different SSP. Relative to current range, seven species showed relative declines in SSP5-8.5, including species that exhibited range increases in less extreme SSP scenarios (Figure 6).

Considering SSP2-4.5 as the high-likelihood climate change scenario, future geographic shifts in species ranges are illustrated in Figure 7. Further southward expansions in SWG are predicted for three species (*Vateria indica*, *Syzygium densiflorum*, *Palaquium ravii*) while others exhibit limited shifts southwards (*Myristica beddomei*, *Orophea thomsonii*, *Diospyros paniculata*). South-westward shifts and contractions of distributions were predicted for three species: *Cryptocarya anamalayana*, *Myristica beddomei*, *Dysoxylum malabaricum*. *Dipterocarpus bourdillonii*, which is highly range-restricted, is predicted to contract further into smaller localised pockets or specific valleys <10 km².

**Figure 7:**
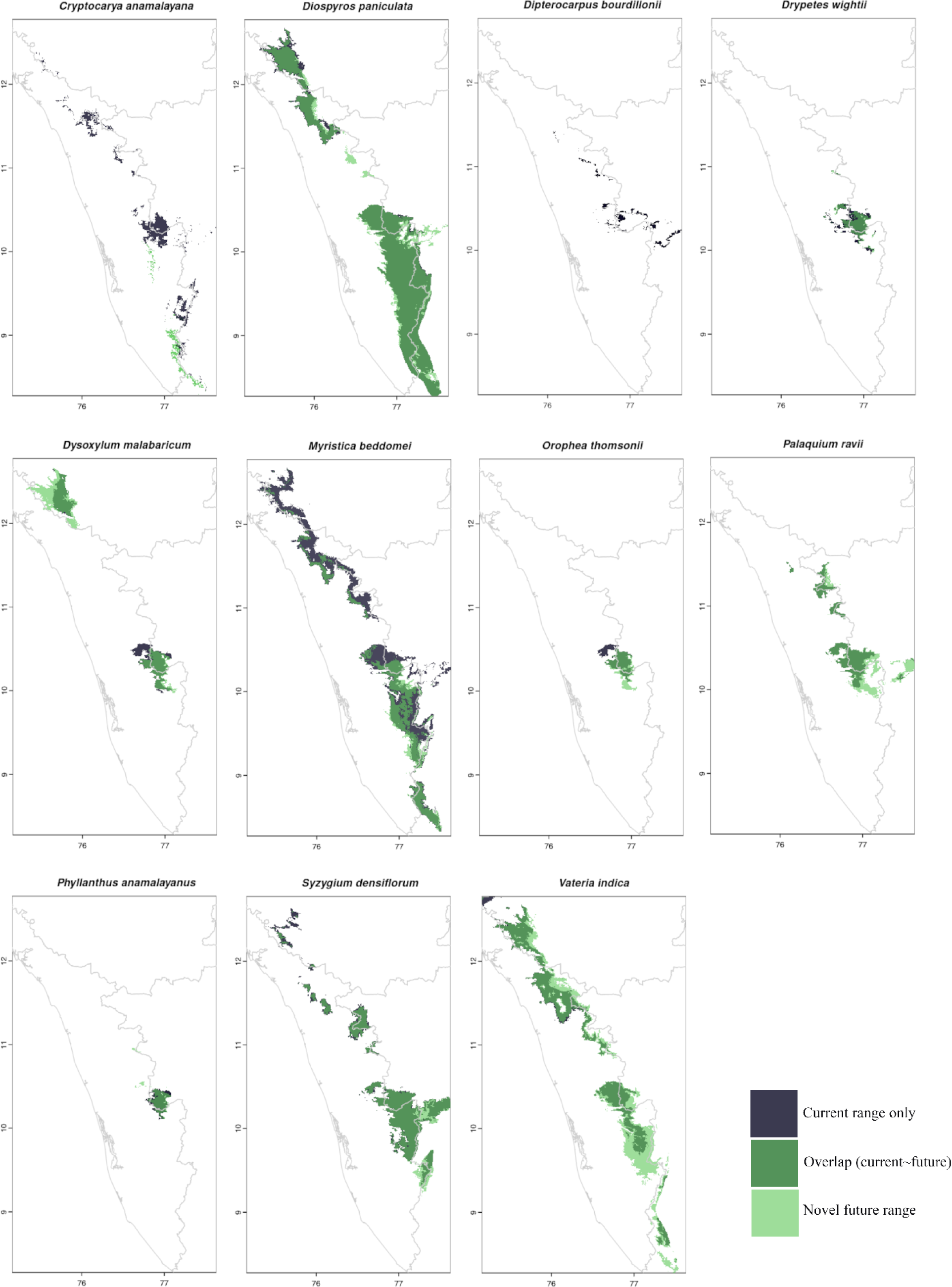
Range changes of the 11 species comparing current range (blue), and predicted future range-overlap(dark green)/ novel (light green) under SSP2-4.5 climate scenario.

## DISCUSSION

The eleven threatened tree species in the present study exhibited a range of responses in abundance in relation to habitat alteration at the landscape level (within the Anamalai Hills) and in distributional ranges in relation to climate at the regional level (SWG). Only the most-widely distributed species (*D. paniculata*) increased in density in rainforest fragments, while 6 species decreased in abundance in fragments (the 7th, *D. bourdillonii* occurred only in the protected area. As the species that declined included both lower- and higher-elevation as well as narrow- and wide-ranging species, the responses likely reflect idiosyncratic and species-specific habitat requirements. While the need for continued conservation of larger rainforest tracts is evident, the rainforest fragments also have conservation value as many threatened trees continue to persist in them, along with a diversity of other endemics and rainforest plant species (Muthuramkumar et al., 2006).

Spatial distributions patterns revealed that 9 of the 11 (excluding *Vateria indica* and *Myristica beddomei*) were restricted to the southern latitudes of the SWG, with 4 species found in highly restricted pockets (*Orophea thomsonii*, *Phyllanthus anamalayanus*, *Palaquium ravii*, *Dipterocarpus bourdillonii*), which is indicative of highly specialised habitat affinities (Belluau & Shipley, 2018; Krishnadas et al., 2016). Species distribution model (SDM) results indicated that precipitation (precipitation of the driest month, precipitation seasonality, precipitation of the wettest quarter) and temperature (mean temperature of the wettest quarter) were the most important influences on current species distributions. Precipitation variables and temperature frequently distinguish occurrence sites from background/reference sites in SDM studies (Bradie & Leung, 2017) and may be particularly significant in ecosystems moulded by precipitation gradients such as the tropical wet evergreen forests of the Southern Western Ghats (SWG) (Krishnadas et al., 2021; Pascal, 1988). Clustering of ranges of habitat specialists in SWG is likely related to past climatic stability resulting in more time for speciation, differentiation and diversification to occur (Bose et al., 2019; Cowling et al., 2015; Harrison & Noss, 2017; Joshi & Karanth, 2013) and the formation of climate refugia acting as ‘museums’ of divergent lineages (Gopal et al., 2023; Vijayakumar et al., 2016).

The influence of precipitation has implications for range shifts following climate change. Within primarily temperature-influenced ecosystems, species ranges are expected to shift to higher elevations under warming climates (Maharjan et al., 2023; McCain & Colwell, 2011). However, within the orographically complex SWG with ecosystems influenced by precipitation gradients, species range shifts may depend on changes in precipitation along combinations of latitude, elevation, seasonality, and aspect (McCain & Colwell, 2011; Tingley et al., 2012). Precipitation-driven range shift models exhibit highly heterogeneous responses based on discordant ‘push’ and ‘pull’ factors of thermoclines (mainly higher-elevation shifts) and precipitation-clines (such as downslope, westward, lower latitudes) (Lenoir et al., 2010; Maharjan et al., 2023; McCain & Colwell, 2011; Tingley et al., 2012). Such processes may thus explain the apparent south-westward shifts in ranges of *Cryptocarya anamalayana*, *Myristica beddomei*, and *Dysoxylum malabaricum*, shifts towards lower latitudes *Syzygium densiflorum*, *Palaquium ravii*, *Orophea thomsonii*, and shifts to lower elevations in *Dysoxylum malabaricum,Vateria indica*. Such trends may interact with biotic factors such as release from competing species or habitat alteration to determine range shifts (Lenoir et al., 2010; Neilson et al., 2005), although this is an aspect requiring further investigation in the SWG.

Climate change projections for the Western Ghats mountain range predict longer, more pronounced seasonal droughts, and increased annual rainfall in the northern latitudes and decrease in southern latitudes, and a greater variation in annual rainfall (Katzenberger et al., 2021). Climate specialists and range-restricted species with narrow environmental tolerances (response curves) are expected to be more sensitive to changes compared to generalists with broader tolerances and greater adaptability and resilience (Bonachela et al., 2021). Our projections indicate varying responses among the threatened trees with some species facing drastic loss or extinction of habitats (‘losers’), while others benefit or remain relatively unchanged (‘winners’) (Dyderski et al., 2018; Garcia et al., 2013; Gilani et al., 2020; Maharjan et al., 2023; Thang et al., 2020). Among both ‘winners’ and ‘losers’ there were lower- and higher-elevation species as well as narrow- and wide-ranging species, again suggesting that climate change responses were individual species-specific rather than directly related to elevational distribution, range size, or degree of specialisation (Godoy-Veiga et al., 2021).

Our models suggest that parts of the SWG will continue to act as important refugia for threatened taxa under future climate change (Chitale et al., 2014; Harrison & Noss, 2017). Existing geographic centres for endemism and biodiversity face severe reductions (∼45%) and relatively cooler more high moisture massifs and valleys (climate refugia) within these hotspots will allow for the persistence of many species communities (Chitale et al., 2014). Locally adapted populations of species and their distances to corresponding climate refugia are important considerations for assessing climatic change risks, species persistence and resilience (Roberts & Hamann, 2016). For the five species for which declines were predicted under all climate scenarios in the present study, specific conservation efforts are needed to conserve the identified refugia where the species are likely to persist into the future. This is particularly relevant for *Dipterocaprus bourdillonii* and *Cryptocarya anamalayana* which already have small populations and narrow ranges. In mountain ecosystems and along elevation gradients, the identification of micro-climatic refugia that enable species to persist under climate change through short distance migration and dispersal is crucial (Harrison & Noss, 2017; Maharjan et al., 2023; Neilson et al., 2005).

Within this study we identify the southern Western Ghats as an ecologically meaningful climate refugia within the context of increasing climate fluctuations. Overall, our results attest the need for conservation of forest tracts within existing protected areas of the SWG while recognising the conservation values of fragments in the wider landscape where many threatened species continue to persist. As responses to climate change are species specific, identifying specific micro-climatically stable zones (refugia) for species populations will be vital and need targeted conservation attention in the future.

## ACKNOWLEDGMENT

We thank the Tamil Nadu Forest Department and the Field Director, Deputy Directors, and Ranger Officers of Anamalai Tiger Reserve for research permits and their cooperation. G. Moorthi, R. Rajesh, and T. Sundarraj assisted in field surveys. We thank Fondation Franklinia, Rohini Nilekani Philanthropies, Rainmatter Foundation, and AMM Murugappa Chettiar Research Centre for funding support to the rainforest research and restoration program.

## Data availability statement

Data for the study are available in Madhavan et al. (2024) Zenodo (Available for reviewers/editors at <u>this link</u>). [The dataset will be made publicly available along with publication.]

