## Supplementary information S1 & S2 for "Distribution models predict climate-related range alteration or extinction of eleven threatened tropical rainforest trees in the Western Ghats"

Figure S1: (next 4 pages) Variable importance and corresponding partial dependence plots for the four most important variables for each of the 11 species.

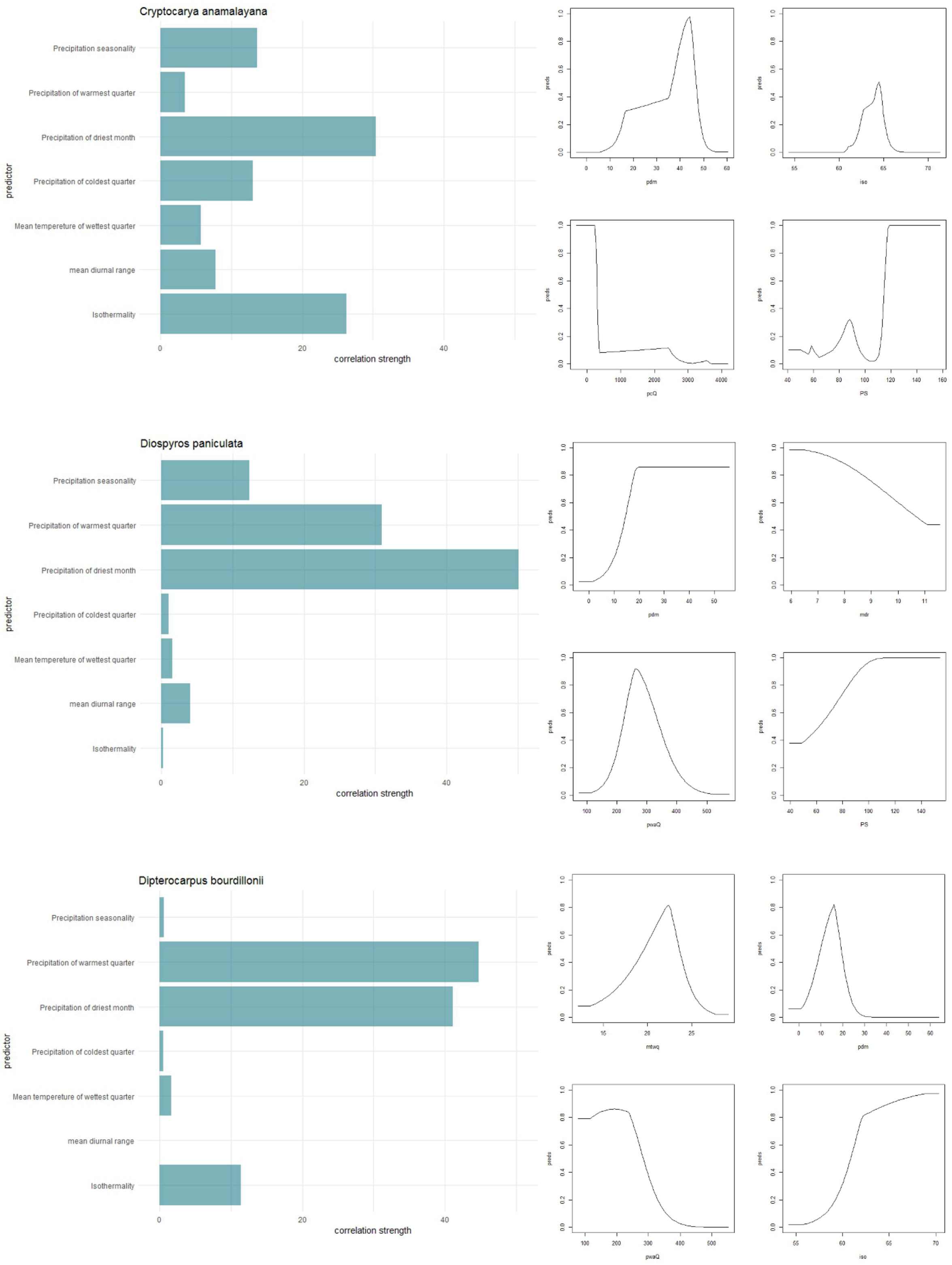

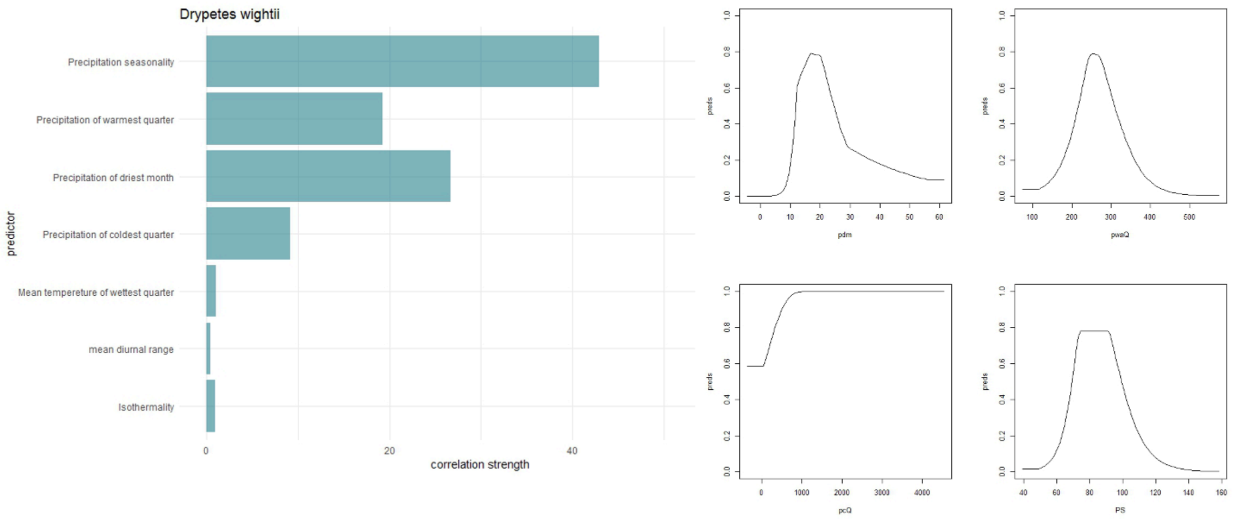

Drypetes wightii
Precipitation seasonality
Precipitation of warmest quarter
Precipitation of driest month
Precipitation of coldest quarter
Mean tempereture of wettest quarter
mean diurnal range
Isothermality
predictor
0
20
40
correlation strength
preds
pdm
pwaQ
pcQ
PS


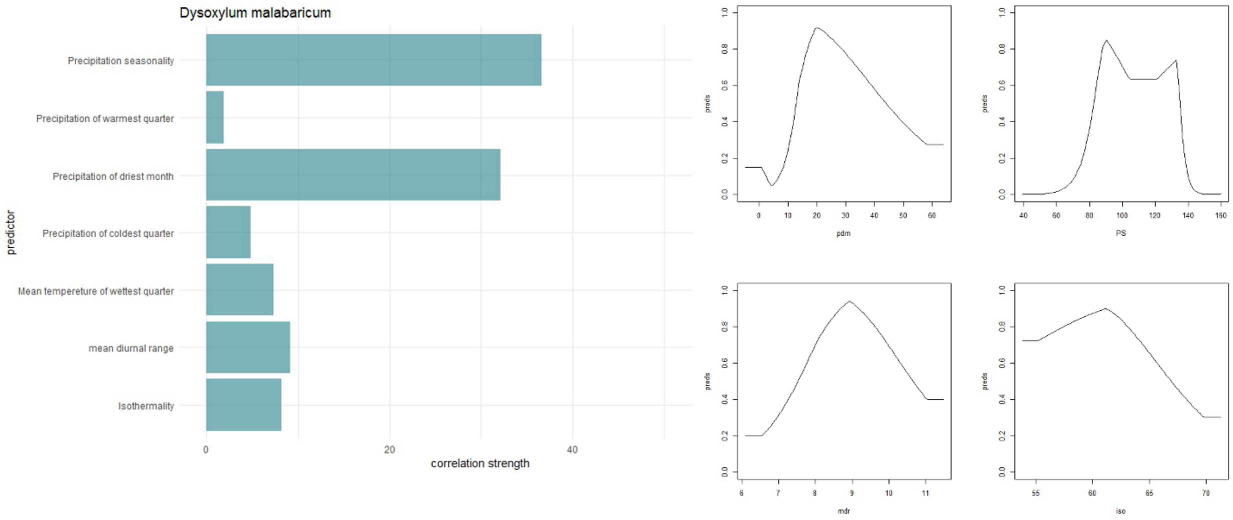

Dysoxylum malabaricum
Precipitation seasonality
Precipitation of warmest quarter
Precipitation of driest month
Precipitation of coldest quarter
Mean tempereture of wettest quarter
mean diurnal range
Isothermality
predictor
0
20
40
correlation strength
preds
pdm
PS
mdr
iso


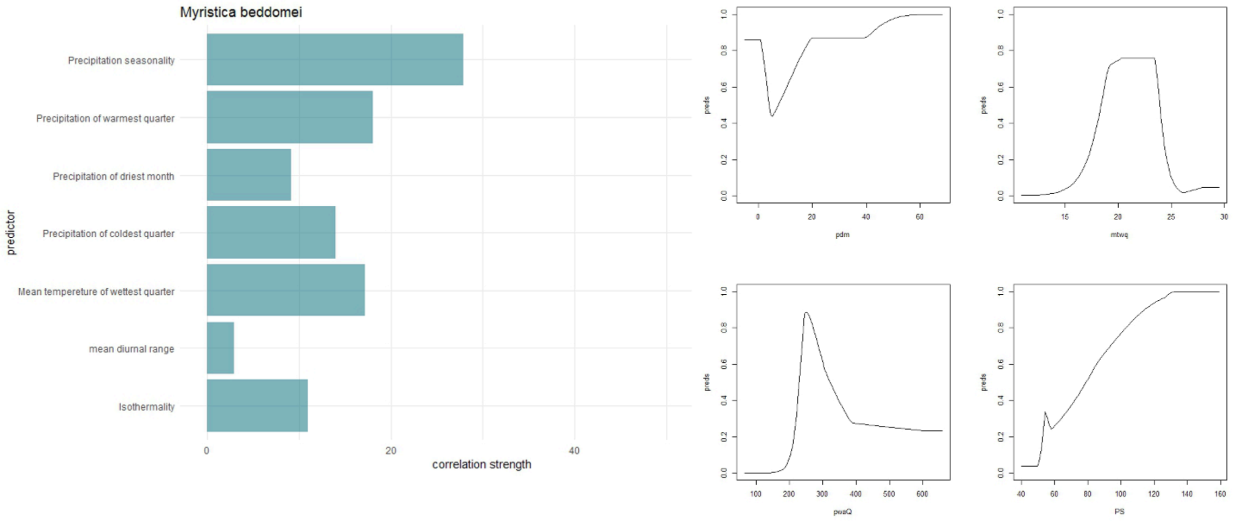

Myristica beddomei
Precipitation seasonality
Precipitation of warmest quarter
Precipitation of driest month
Precipitation of coldest quarter
Mean tempereture of wettest quarter
mean diurnal range
Isothermality
predictor
0
20
40
correlation strength
preds
pdm
mtwq
pwaQ
PS

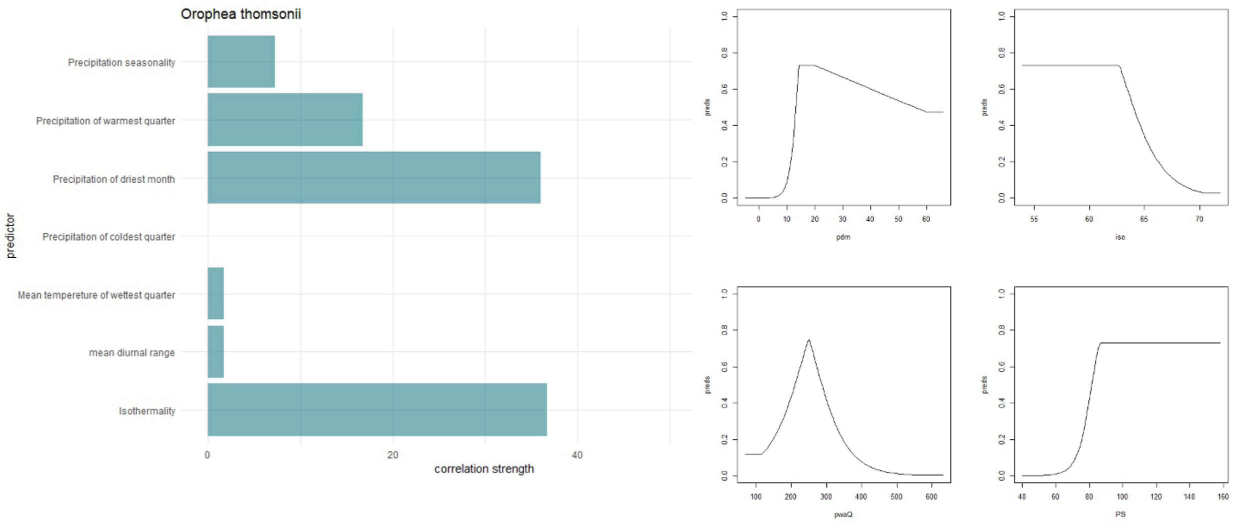
Orophea thomsonii
Precipitation seasonality
Precipitation of warmest quarter
Precipitation of driest month
Precipitation of coldest quarter
Mean tempereture of wettest quarter
mean diurnal range
Isothermality
predictor
0
20
40
correlation strength
preds
pdm
iso
pwaQ
PS

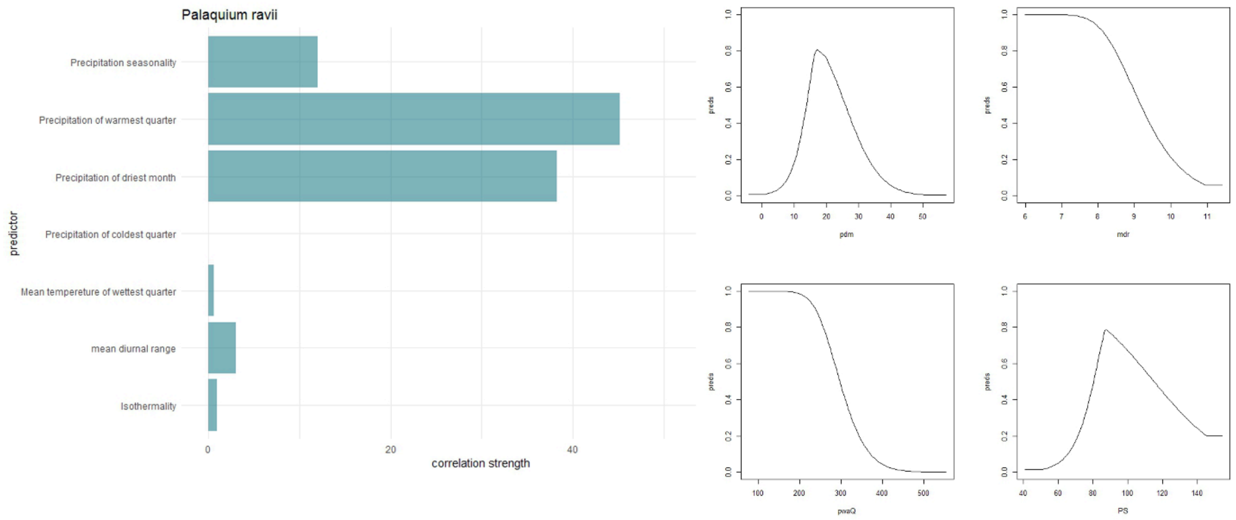
Palaquium ravii
Precipitation seasonality
Precipitation of warmest quarter
Precipitation of driest month
Precipitation of coldest quarter
Mean tempereture of wettest quarter
mean diurnal range
Isothermality
predictor
0
20
40
correlation strength
preds
pdm
mdr
pwaQ
PS

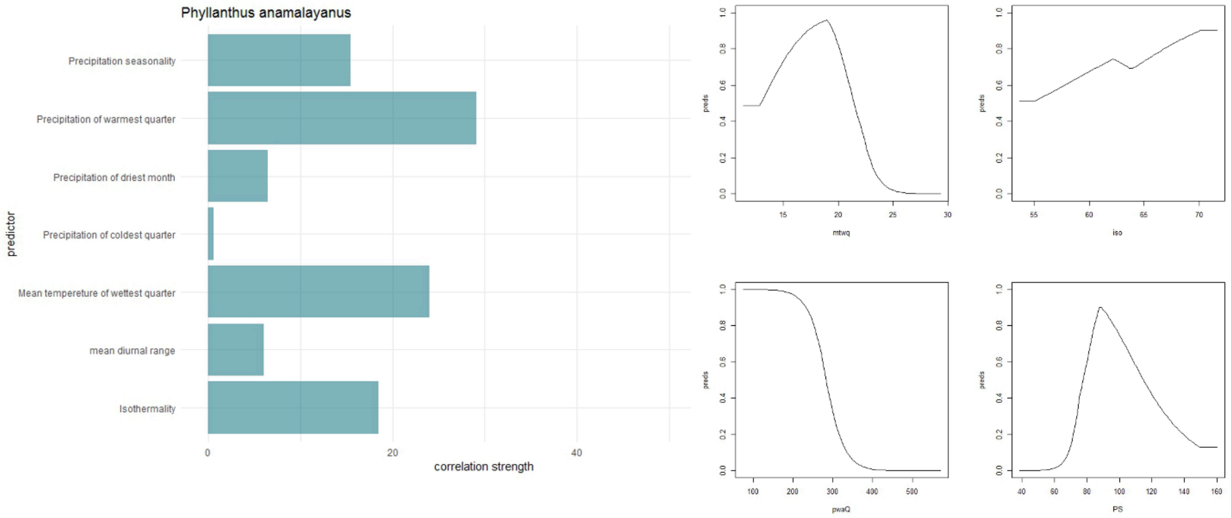
Phyllanthus anamalayanus
Precipitation seasonality
Precipitation of warmest quarter
Precipitation of driest month
Precipitation of coldest quarter
Mean tempereture of wettest quarter
mean diurnal range
Isothermality
predictor
0
20
40
correlation strength
preds
mtwq
iso
pwaQ
PS

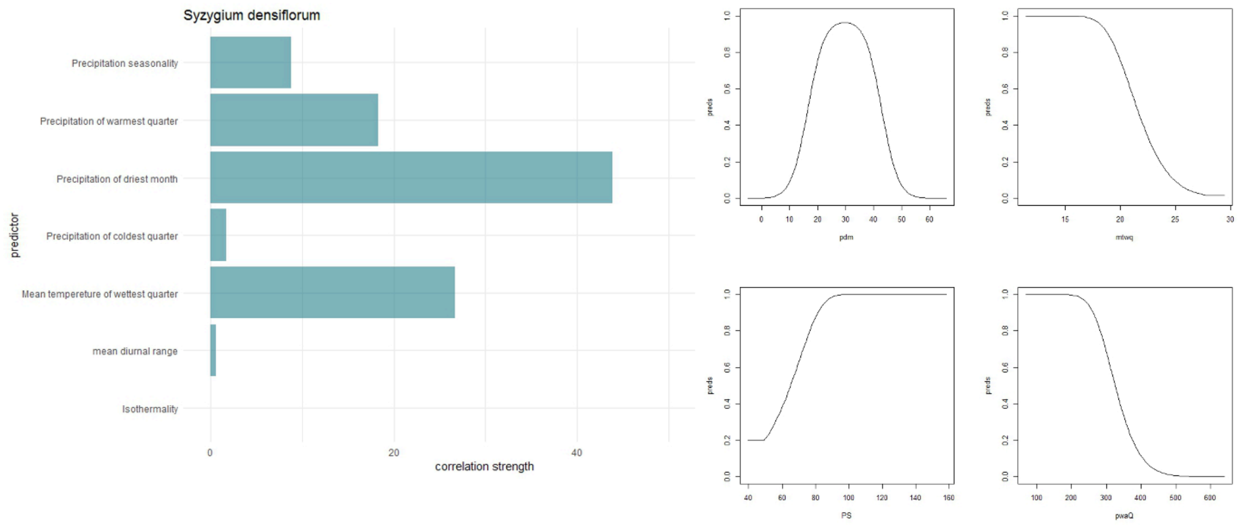
Syzygium densiflorum
Precipitation seasonality
Precipitation of warmest quarter
Precipitation of driest month
Precipitation of coldest quarter
Mean tempereture of wettest quarter
mean diurnal range
Isothermality
predictor
0
20
40
correlation strength
preds
pdm
mtwq
PS
pwaQ

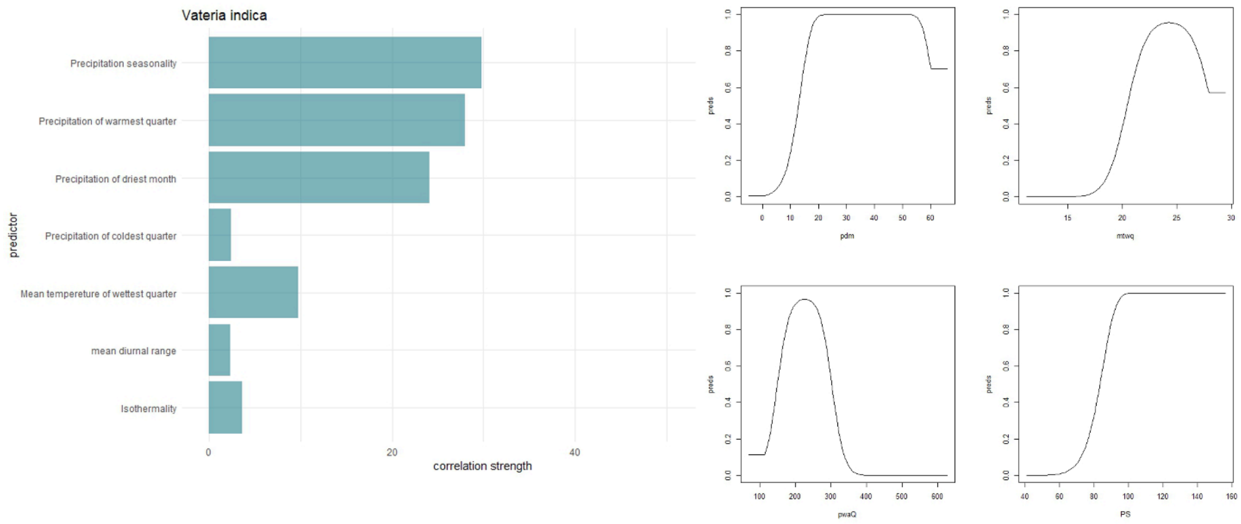
Vateria indica
Precipitation seasonality
Precipitation of warmest quarter
Precipitation of driest month
Precipitation of coldest quarter
Mean tempereture of wettest quarter
mean diurnal range
Isothermality
predictor
0
20
40
correlation strength
preds
pdm
mtwq
pwaQ
PS

Figure S2: Niche breadth (Levins B2) value for the eleven species

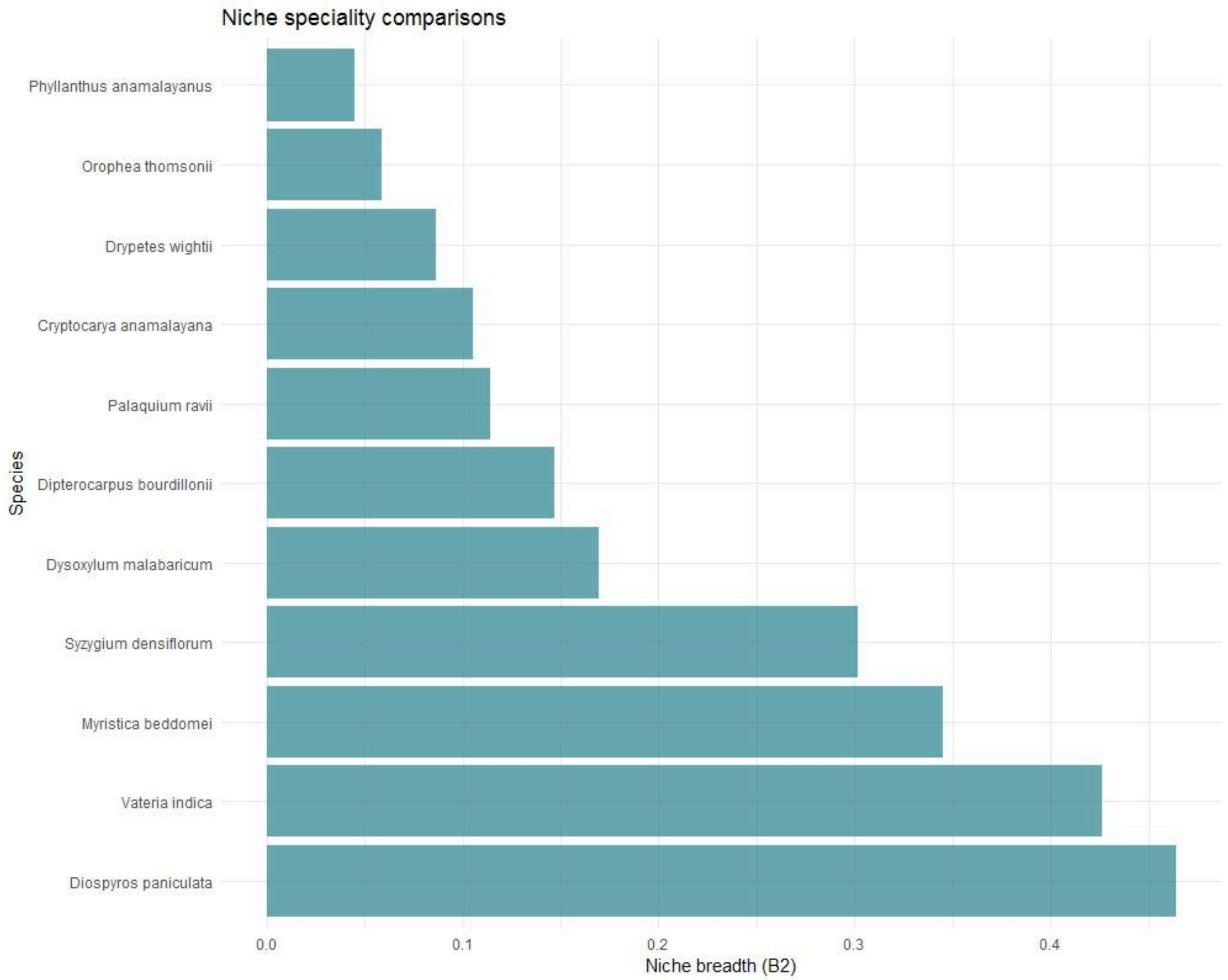
